# Global profiling of the RNA and protein complexes of *Escherichia coli* by size exclusion chromatography followed by RNA sequencing and mass spectrometry (SEC-seq)

**DOI:** 10.1101/2022.09.02.506378

**Authors:** Kotaro Chihara, Milan Gerovac, Jens Hör, Jörg Vogel

## Abstract

New methods for the global identification of RNA-protein interactions have led to greater recognition of the abundance and importance of RNA-binding proteins (RBPs) in bacteria. Here, we expand this tool kit by developing SEC-seq, a method based on a similar concept as the established Grad-seq approach. In Grad-seq, cellular RNA and protein complexes of a bacterium of interest are separated in a glycerol gradient, followed by high-throughput RNA-sequencing and mass spectrometry analyses of individual gradient fractions. New RNA-protein complexes are predicted based on the similarity of their elution profiles. In SEC-seq, we have replaced the glycerol gradient with separation by size exclusion chromatography, which shortens operation times and offers greater potential for automation. Applying SEC-seq to *Escherichia coli*, we find that the method provides a higher resolution than Grad-seq in the lower molecular weight range up to ∼500 kDa. This is illustrated by the ability of SEC-seq to resolve two distinct, but similarly sized complexes of the global translational repressor CsrA with either of its antagonistic small RNAs, CsrB and CsrC. We also characterized changes in the SEC-seq profiles of the small RNA MicA upon deletion of its RNA chaperones Hfq and ProQ and investigated the redistribution of these two proteins upon RNase treatment. Overall, we demonstrate that SEC-seq is a tractable and reproducible method for the global profiling of bacterial RNA-protein complexes that offers the potential to discover yet-unrecognized associations between bacterial RNAs and proteins.

## INTRODUCTION

Most biological processes depend on stable cellular complexes. These often include RNAs, which together with proteins, form higher-order ribonucleoprotein particles (RNPs). In bacteria, several RNA-binding proteins (RBPs) function as structural components of RNPs (Gerovac et al. 2021a). These stable RNPs range in size from the 70 kDa signal recognition particle (SRP) involved in the co-translational translocation of proteins to the membrane (Akopian et al. 2013) to the giant 70S ribosome assembled by three ribosomal RNA species and >50 ribosomal proteins (Davis and Williamson 2017).

In addition to their function as structural RNP components, bacterial RBPs can also act as post-transcriptional regulators of different classes of RNAs (Holmqvist and Vogel 2018; Ng Kwan Lim et al. 2021; Christopoulou and Granneman 2022). For example, Hfq, one of the best-known bacterial RBPs, functions as an RNA chaperone in both gram-negative and gram-positive bacteria (Vogel and Luisi 2011; Kavita et al. 2018). Hfq helps small RNAs (sRNAs) base-pair with target mRNAs (Møller et al. 2002; Zhang et al. 2002). Thereby, it contributes to translational repression or activation of these mRNAs and their subsequent stabilization or degradation, respectively (Morita and Aiba 2011; De Lay et al. 2013; Schu et al. 2015). The ProQ/FinO-domain proteins are an emerging new family of global post-transcriptional regulators in gram-negative bacteria (Attaiech et al. 2016; Smirnov et al. 2016; Olejniczak and Storz 2017; Holmqvist et al. 2020). ProQ binds to sRNAs and also appears to be responsible for the base-paring between sRNAs and target mRNAs and subsequent post-transcriptional regulation (Smirnov et al. 2017b; Melamed et al. 2019). In contrast, CsrA, another well-established RBP, primarily binds to the 5’UTRs of target mRNAs directly and affects translation by occluding or promoting ribosome association (Potts et al. 2017; Pourciau et al. 2020). Post-transcriptional regulation by CsrA is inhibited by the sRNAs CsrB and CsrC, which antagonize the interaction of CsrA with its mRNA targets (Dubey et al. 2005; Duss et al. 2014). Despite the functional importance of these established RBPs, they interact with only a small subset of bacterial RNAs. Many additional putative bacterial RBPs have been predicted bioinformatically (Sharan et al. 2017), and the recent increase in the number and diversity of CRISPR-Cas systems has further contributed to the discovery of structurally and functionally diverse bacterial RNPs (van der Oost et al. 2014; Makarova et al. 2019). It therefore seems certain that additional RBPs await discovery.

Experimental investigations of RNA-protein complexes with newly developed methods have been extensively carried out in eukaryotes (Hentze et al. 2018), but these methods developed for eukaryotic cells are not directly transferable to bacteria, whose transcripts lack a functional poly(A) tail and are resistant to artificial nucleotides. While several global approaches that overcome these challenges have recently been developed, these methods rely mostly on UV crosslinking of components of an RNA-protein complex, followed by either organic phase extraction or silica-based solid-phase purification (Asencio et al. 2018; Trendel et al. 2018; Queiroz et al. 2019). Although these methods represent powerful tools that have been applied successfully to enteric bacteria to enrich Hfq, ProQ, and additional RBP candidates (Shchepachev et al. 2019; Urdaneta et al. 2019; Chu et al. 2022), UV crosslinking biases the recovery of RBPs to those containing pyrimidines, especially uracil, in their binding sites (Wheeler et al. 2018). Hence, there remains a need for method development for the discovery and study of bacterial RBPs.

We have recently introduced Grad-seq, an unbiased gradient sequencing approach to predict new protein-RNA complexes in bacteria (Smirnov et al. 2016; Smirnov et al. 2017a). In Grad-seq, native cell lysates are loaded onto a linear glycerol gradient and subjected to ultracentrifugation. All soluble molecules are separated into fractions depending on molecular weight. RNAs and proteins are purified from each fraction and analyzed by high-throughput RNA-sequencing (RNA-seq) and mass spectrometry (MS). Genome-wide reconstitution of in-gradient distribution of all detectable RNAs and proteins allows the prediction of RBPs and their RNA targets (Smirnov et al. 2017a). Applying this approach to various bacterial species has contributed to the discovery of the RNA chaperone ProQ in *Salmonella enterica* serovar Typhimurium (henceforth *Salmonella*) (Smirnov et al. 2016), of the previously unrecognized association of sRNAs and a small protein with the *Escherichia coli* ribosome (Hör et al. 2020a), and of novel RBPs in gram-positive bacteria and in cyanobacteria (Hör et al. 2020b; Riediger et al. 2020; Lamm-Schmidt et al. 2021). Grad-seq profiles of *Pseudomonas aeruginosa* after bacteriophage infection have highlighted the capacity of this method to reveal the global reprogramming of RNA-protein complexes (Gerovac et al. 2021b). Overall, Grad-seq has provided widely-used resources of RNA-protein complexes for both gram-negative and gram-positive bacteria. In addition, Grad-seq has been successfully applied to eukaryotic cells (Aznaourova et al. 2020; Schneider et al. 2022). However, the approach requires a long operation time for ultra-centrifugation and has only limited potential for automation.

Here, we develop an alternative approach to Grad-seq, based on the same principles and readouts, but using size exclusion chromatography (SEC) for the separation of cell lysates. SEC is a powerful tool to resolve RNAs and proteins depending on their size and shape (Kirkwood et al. 2013; Wan et al. 2015; Larance et al. 2016; Yoshikawa et al. 2018; Mallam et al. 2019; Skinnider et al. 2021). SEC fractionation requires a shorter operation time (2 h for SEC gel-filtration in SEC-seq vs. 17 h for glycerol gradient centrifugation in Grad-seq), which is likely to reduce RNA degradation.

Furthermore, eluates are automatically separated so that laborious manual fractionation is not required. For proof-of-concept, we used the method to resolve the profiles of ∼85% of all transcripts and ∼60% of proteins of *E. coli*. Importantly, we observed a higher resolution in the size range of less than ∼500 kDa compared to Grad-Seq, which is illustrated by the distinct elution profiles of CsrA-CsrB and CsrA-CsrC, two similarly sized complexes of CsrA with either of its antagonistic small RNAs. Thus, we show that SEC-seq is a tractable and reproducible method for RNA-protein complex profiling. The SEC-seq data generated in this study have been integrated into an online browser available at https://helmholtz-hiri.de/en/datasets/gradseqec/ that allows users to cross-compare the distributions of RNAs or proteins of interest between Grad-seq and SEC-seq (Hör et al. 2020a).

## RESULTS AND DISCUSSION

### SEC allows fractionation of cellular RNAs and proteins in gram-negative and gram-positive bacteria

To set up the SEC-seq approach, we separated RNAs and proteins of an *E. coli* native lysate obtained from an early stationary phase (OD_600_ = 2.0) through a Superose 6 Increase SEC column (Fig. 1A). The general SEC chromatogram shows four major peaks (Fig. 1B). Total RNA and protein were then extracted from these peaks and analyzed by Urea-PAGE or SDS-PAGE, respectively (Supplemental Fig. S1). We observed few RNAs and proteins in peak 4 compared to peaks 1-3, suggesting that peak 4 mainly includes degradation products and proteins or peptides smaller than 10 kDa (Supplemental Fig. S1B–C). Based on the known fractionation range of the Superose 6 Increase SEC column (5 to 5000 kDa), peak 1 likely consists of aggregates (Fig. 1B; Supplemental Fig. S1A). Therefore, we focused on the eluate ranging from peak 2 to peak 3, separated into 20 fractions (elution volume = 0.38 ml). This range also corresponds to the range of glycerol gradients (Smirnov et al. 2016; Hör et al. 2020a). For easier comparison with a previously performed *E. coli* Grad-seq experiment (Hör et al. 2020a), we numbered fractions 1-20 starting from low molecular weight (LMW) to high molecular weight (HMW).

**Figure 1.**
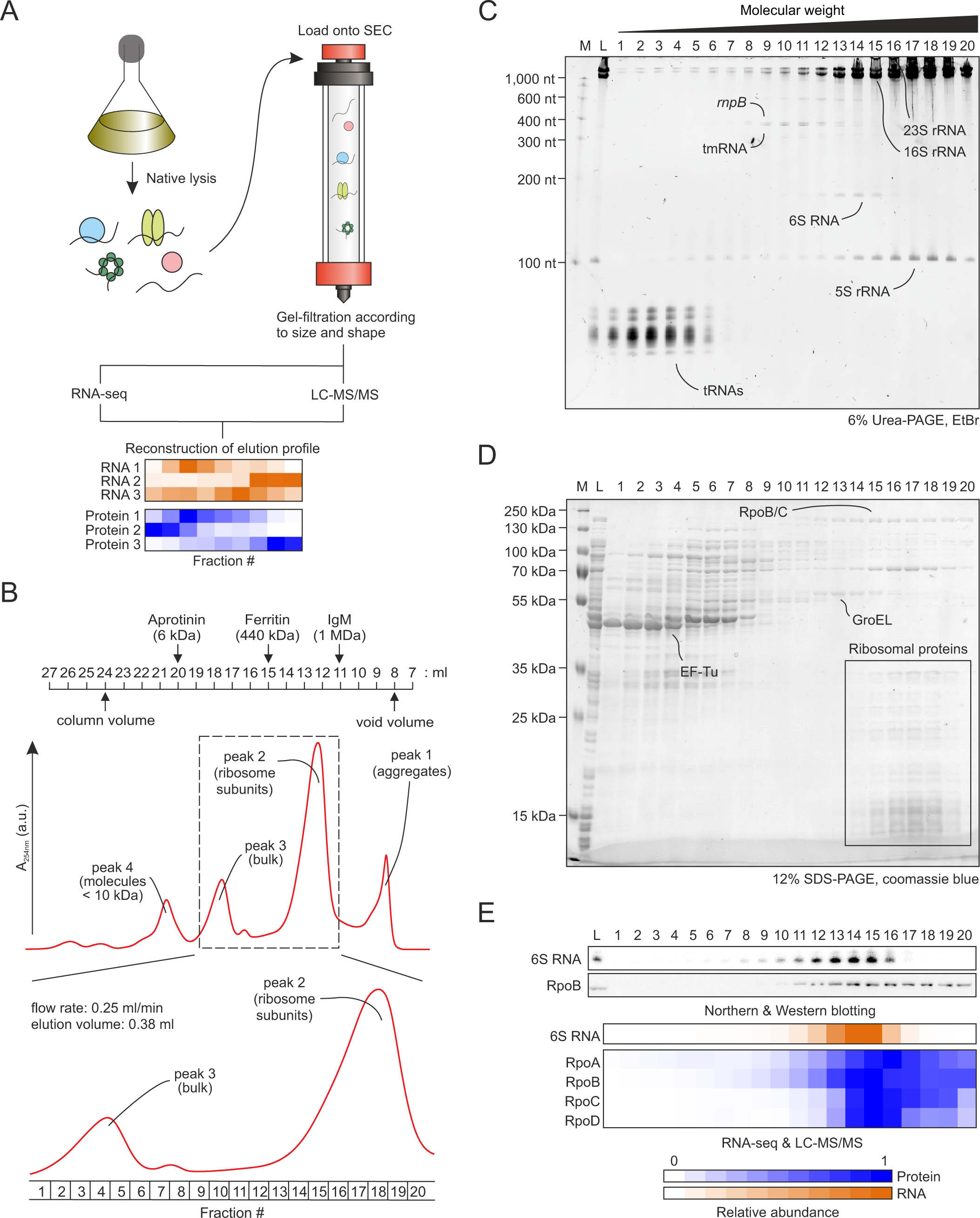
Separation and analysis of the *E. coli* lysate by SEC. (A) Overview of the SEC-seq approach. The native cell lysate is loaded onto tshe SEC column and fractioned based on the size and shape of the particles. The elution profiles of all RNAs and proteins detected by RNA-seq and LC-MS/MS are reconstituted. (B) SEC chromatogram shows four major peaks and the region between peak 2 and peak 3 is further investigated. Elution volume is set to 0.38 ml. Size markers were added from the manual of Superose 6 Increase 10/300 GL (Cytiva 2020). (C and D) Visualization of major RNA (C) and protein (D) elution profiles by Urea-PAGE and SDS-PAGE. High abundant house-keeping RNAs and proteins are indicated. (E) Validation of RNA-seq and MS using conventional northern and western blotting analyses. 6S RNA was co-eluted with RNAP subunits RpoB. L, lysate; M, marker.

Urea-PAGE and SDS-PAGE showed the expected elution profiles of known RNAs and their protein partners (Fig. 1C–D). For example, in HMW fractions (fractions 14–20), rRNAs co-eluted with ribosomal proteins. In contrast, tRNAs co-localized with elongation factor Tu (EF-Tu) in LMW fractions, as previously observed in Grad-seq profiles of *E. coli*, *Salmonella,* or *P. aeruginosa* (Smirnov et al. 2016; Hör et al. 2020a; Gerovac et al. 2021b). We also assessed the distribution of 6S RNA and its well-known interaction partner, the RNA polymerase (RNAP) subunit RpoB (Wassarman 2018) by northern and western blot analysis, respectively. As expected, we observed correlated profiles of 6S RNA and RpoB in fractions 11 to 16 (Fig. 1C–E). To obtain a global view of the distributions of RNAs and proteins separated by SEC, purified RNAs and proteins from fractions 1–20 were subjected to RNA-seq and MS analyses, respectively (Fig. 1A). These profiles confirmed the distribution of 6S RNA and RpoB, as well as other RNAP subunits (Fig. 1E), and will be described in more detail below.

To determine if SEC fractionation can be expanded to other bacterial species, we tested lysates from the gram-negative bacterium *P. aeruginosa* and gram-positive *Clostridioides difficile* (Supplemental Fig. S2). In both species, rRNAs co-eluted with the 50S and 30S ribosomal subunits in HMW fractions, while tRNAs co-localized with EF-Tu in LMW fractions (Supplemental Fig. S2A–B). The overall profiles of representative RNAs and proteins for both bacterial species were similar to *E. coli*, confirming that SEC fractionation can be applied to both gram-negative and gram-positive bacteria.

### Global profiling of RBPs and protein complexes by SEC-seq

Next, we examined the overall distribution of *E. coli* RNAs and proteins along the SEC column. Based on the annotations from RefSeq (Pruitt et al. 2007), RegulonDB (Huerta et al. 1998), and ANNOgesic (Yu et al. 2018), RNA-seq of the 20 SEC fractions captured the profiles of 4,194 transcripts (using a threshold of > 100 reads in the sum of fractions), comprising 3,787 mRNAs, 298 sRNAs, and all tRNAs and rRNAs (Supplemental Fig. S3; Supplemental Table S1). MS analysis of the same fractions reported SEC profiles of 2,428 proteins using a threshold of > 2 identified razor/unique peptides based on the annotation from UniProt (UniProt 2019) (Supplemental Table S2). Since we used cleared cellular lysate, we recovered primarily cytosolic proteins (60% of all detected proteins) (Supplemental Fig. S4). Relative abundance of all detected transcripts and proteins in each fraction was reproducible between the two replicates (Supplemental Fig. S5A–B). The individual SEC profiles of transcripts and proteins also correlated well in both replicates (Supplemental Fig. S5C–D), indicating that SEC-seq provides reproducible results.

Focusing on known RBPs and protein complexes, we observed that these RBPs generally eluted in similar fractions as their established RNA or protein partners (Fig. 2). For instance, ribosomal proteins eluted in fractions 14 to 20, as did rRNAs (Fig. 1C). Ribosome-associated proteins and rRNA modification factors, on the other hand, populated distinct fractions, suggesting their condition-dependent or transient association with ribosomes. To illustrate, the 23S rRNA m^1^G methyltransferase RlmA, which interacts with free 23S rRNA but not assembled ribosomes, eluted in LMW fractions. Whereas 23S rRNA 2’-*O*-ribose methyltransferase RlmE, which is active on ribosomes but not free 23S rRNA, eluted in HMW fractions (Caldas et al. 2000; Hansen et al. 2001). Although most amino-acyl-tRNA synthetases and tRNA modification factors eluted in fractions 3 to 7, we observed AlaS, PheST, and SelAB, which form large multimers (AlaS ≈ 250 kDa, PheST ≈ 250 kDa, and SelAB ≈ 500 kDa), in HMW fractions (Fig. 2) (Fayat et al. 1974; Dignam et al. 2011; Manzine et al. 2013). We also noticed that five RNases known to cleave pre-tRNAs (Rbn: RNase BN; Rnb: RNase II; Rnd: RNase D; Rnt: RNase T; Rph: truncated RNase PH) all eluted in similar fractions as tRNAs (Kitamura et al. 1977; Cudny and Deutscher 1980; Deutscher et al. 1988; Reuven and Deutscher 1993; Dutta et al. 2013; Czech 2020). In contrast, RNases involved in rRNA precursor cleavage (Rna: RNase I; YciV: RNase AM) eluted in ribosomal fractions (Kaplan and Apirion 1975; Jain 2020). RNases involved in mRNA turnover (Rng: RNase G; Rnr: RNase R; Pnp: polynucleotide phosphorylase, a.k.a PNPase) showed broad distribution in fractions 8–20, consistent with the distribution of mRNAs. However, RNase E (Rne), the major endoribonuclease for mRNA turnover, elutes exclusively in fractions 18–20, consistent with RNase E forming a multiprotein complex ‘degradosome’ with the RhlB RNA helicase, PNPase, and enolase, as previously reported (Py et al. 1994; Py et al. 1996). While RhlB indeed co-localized with RNase E, PNPase and enolase did not elute in the same fractions as RNase E (Supplemental Table S2). This is in line with an earlier study, which showed that PNPase and enolase are present in large excess compared to RNase E and RhlB, and therefore predominantly exist independent of the degradosome (Liou et al. 2001). Overall, these protein distributions validate SEC-seq as a suitable method to separate bacterial RNA and protein complexes.

**Figure 2.**
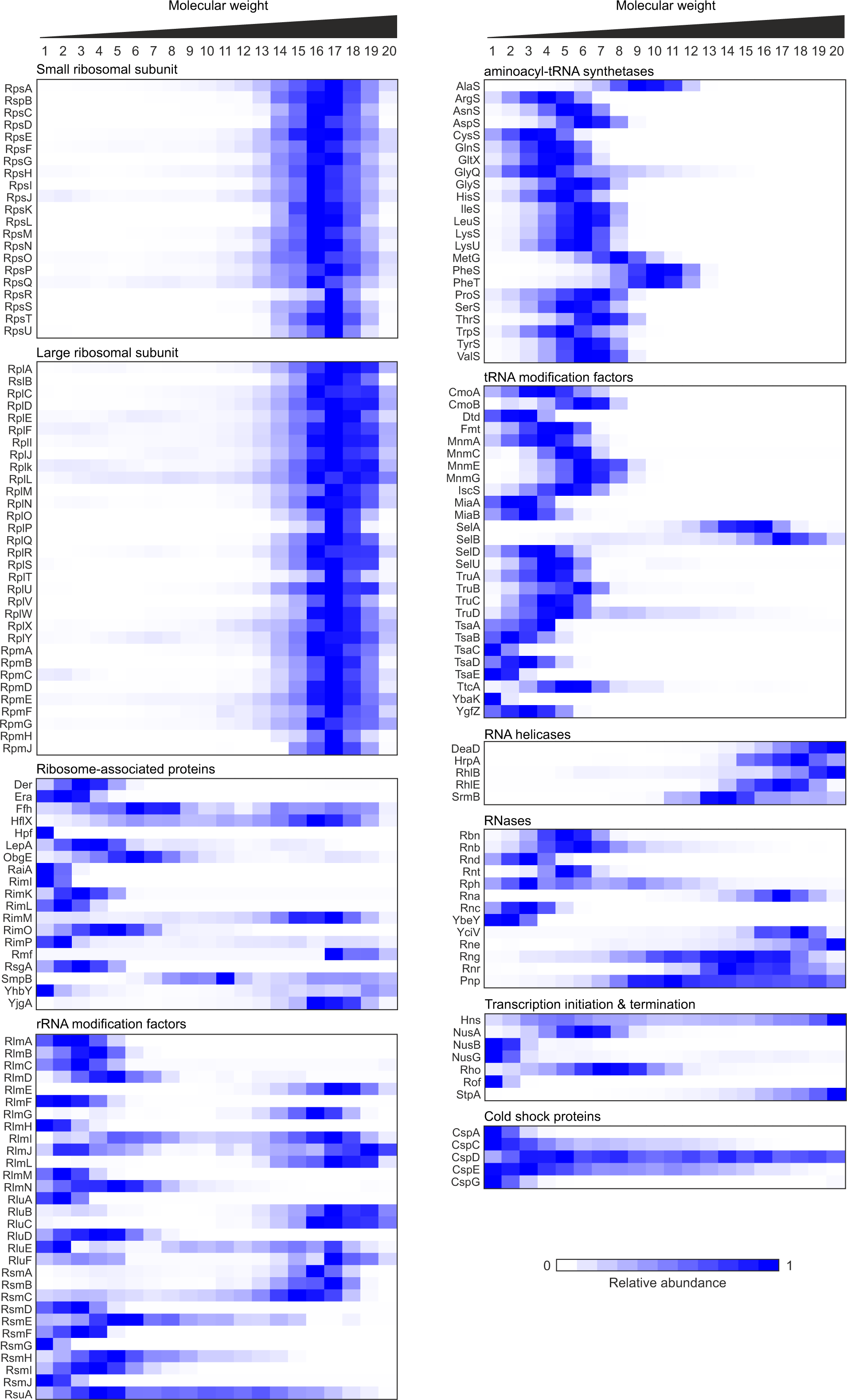
Overview of the elution profiles of known RBPs and the associated proteins. Heat map showing the elution profiles of detected RBPs. They were generally eluted in similar fractions to their specific target RNA or protein partners. Ribosome- associated proteins and rRNA modification factors are distributed through the fractions. For each protein, the spike-in-normalized elution profiles are normalized to the range from 0 to 1 by dividing the values of each fraction by the maximum value of the corresponding protein.

Next, we explored less well-studied RBPs and focused on cold shock proteins (CSPs). *E. coli* encodes 9 CSPs, termed CspA to CspI, which have been reported to interact with hundreds of transcripts, albeit with low affinity (Phadtare and Inouye 1999; Michaux et al. 2017; Yair et al. 2022). In our SEC-seq experiment, we detected CspA, CspC, CspD, CspE, and CspG, which mostly eluted in the first LMW fractions (10-20 kDa), suggesting transient interactions with target transcripts. Only CspD was present in a large range of fractions, implying that CspD forms stable complexes with either other proteins or RNAs. Intriguingly, CspD is an exceptionally toxic protein that inhibits DNA replication (Yamanaka and Inouye 1997; Yamanaka et al. 2001), suggesting that it might require tight regulation through RNA or protein partners. This example highlights the power of SEC-seq to identify interesting candidates for new cellular complexes that warrant further investigation.

### SEC-seq shows improved resolution in the LMW range compared to Grad-seq

Given the different physical principles of particle separation underlying SEC-seq and Grad-seq, we were interested in comparing the resolution of both methods. We first plotted the reported sedimentation coefficients of molecular complexes and their stokes radius, which is defined by particle size and shape (Siegel and Monty 1966; Erickson 2009), against their peak fractions in Grad-seq or SEC-seq, respectively (Fig. 3A). As expected, the sedimentation coefficient and the stokes radius are proportional to the peak fraction in Grad-seq and SEC-seq, respectively. This confirms that SEC fractionation depends linearly on the size and shape of the molecules (Erickson 2009).

**Figure 3.**
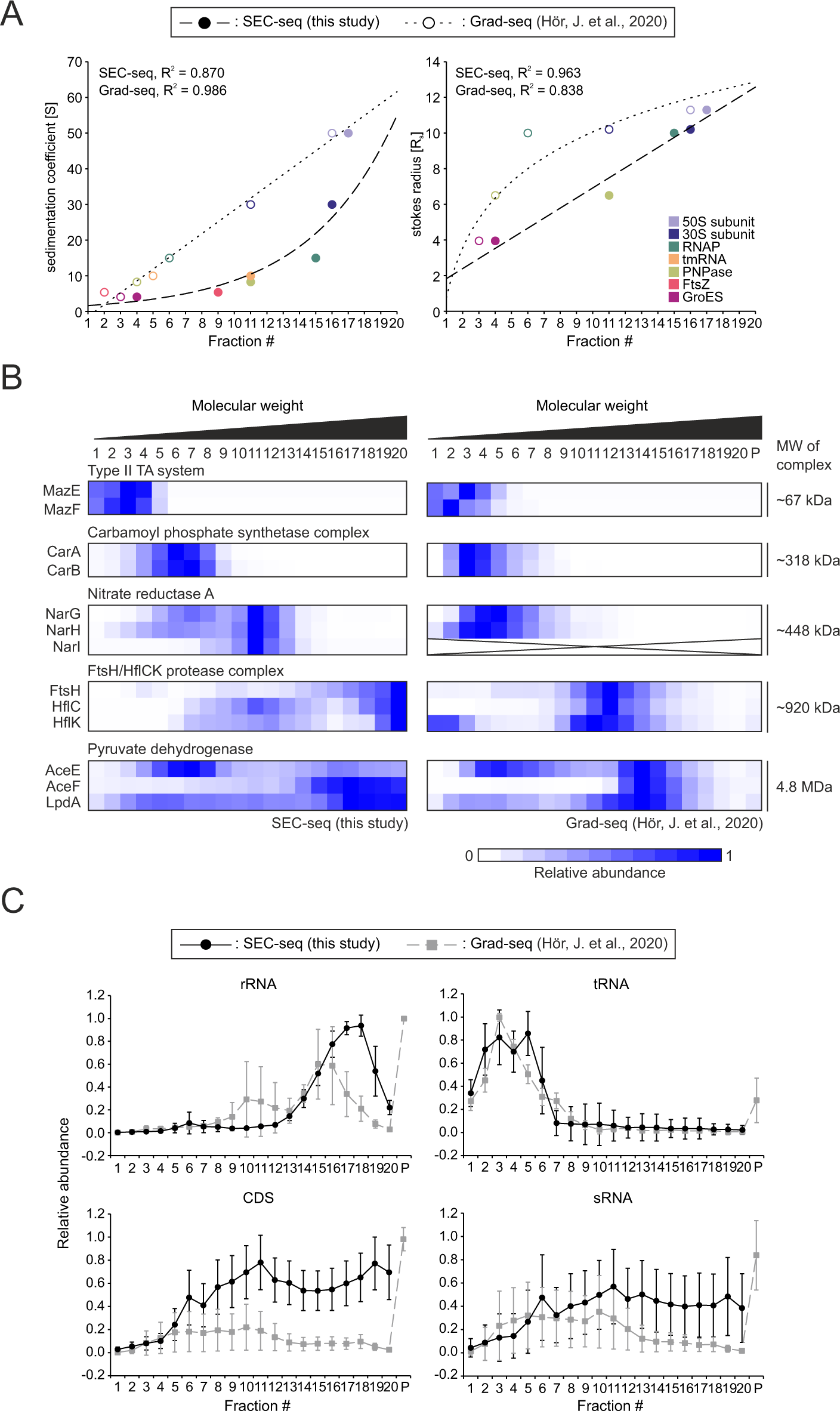
Comparison of RNA and protein profiles between SEC-seq and Grad-seq. (A) The sedimentation coefficient (left) and the stokes radius (right) of molecular particles plotted against their peak fractions obtained from SEC-seq (close circles) and Grad-seq (open circles). The dashed and dotted lines are fitted curves for SEC-seq and Grad-seq, respectively. R^2^ is the coefficient of determination. FtsZ: 5.4 [S] (Rivas et al. 2000); GroES: 4.1 [S], 3.95 [R_s_] (Chandrasekhar et al. 1986; Seale et al. 1996);PNPase: 8.3 [S], 6.5 [R_s_] (Portier 1975; Modrak-Wójcik et al. 2007); tmRNA: 10 [S] (Ray and Apirion 1979); RNAP: 15 [S], 10 [R_s_] (Iwakura et al. 1974; Austin et al. 1983); 30S subunit: 30 [S], 10.2 [R_s_] (Gabler et al. 1974); 50S subunit: 50 [S], 11.3 [R_s_] (Gabler et al. 1974). (B) Comparison of typical protein-protein complexes between SEC-seq and Grad-seq. Complexes spanning from 60 kDa to 450 kDa are well resolved in SEC-seq, whereas larger complexes show more distinct allocation in Grad-seq. The data is scaled to the maximum value. NarI was not detected in Grad-seq. (C) Averaged SEC-seq and Grad-seq profiles of major RNA classes; rRNA, tRNA, mRNA, and sRNA. All individual profiles of RNAs from each class are presented as an average along the fraction ±SD. The rRNA profile from Grad-seq shows two major peaks representing 30S and 50S ribosome subunits while the SEC-seq profile shows one major peak. The SEC-seq profiles of mRNAs and sRNAs are more heterogeneous than Grad-seq.

SEC-seq resolved complexes from 10 kDa (GroES) to ∼500 kDa (the RNAP holoenzyme) in fractions 4 to 15, while in Grad-seq the RNAP holoenzyme and the 1.5 MDa 50S ribosomal subunits eluted from fractions 6 to 16. This indicates that SEC-seq shows a greater resolution in the LMW range up to 500 kDa compared to Grad-seq, while Grad-seq has a greater resolution in the HMW range. To confirm this trend, we compared the SEC-seq and Grad-seq profiles of several protein complexes, ranging from a small 70 kDa type II toxin-antitoxin system to the large 4.8 MDa pyruvate dehydrogenase complex (Fig. 3B). Indeed, small- to mid-sized complexes in the 50 to 500 kDa range, including the toxin-antitoxin complex MazEF, the carbamoyl phosphate synthetase complex, and nitrate reductase A, were better resolved across fractions 1 to 13 in SEC-seq, while they all co-migrated in fraction 1 to 7 in Grad-seq (Fig. 3B). These examples reiterate the improved LMW range resolution of SEC-seq compared to Grad-seq.

Next, we compared the average distributions of transcripts classified into different RNA classes (rRNA, tRNA, CDS, and sRNA) between SEC-seq and Grad-seq (Fig. 3C). Additionally, we calculated the relative positions of each RNA class in SEC-seq and Grad-seq profiles as the mean of proportions of transcripts times each fraction number (see METHOD). This allows us to evaluate whether the difference in the overall profiles between SEC-seq and Grad-seq is significant or not (Supplemental Fig. S6A-D). Grad-seq showed two major peaks derived from 16S and 5S/23S rRNAs, respectively, whereas SEC-seq showed the co-elution of 16S and 5S/23S rRNAs in fractions 14–20, affirming that Grad-seq has an advantage over SEC-seq for the separation of HMW particles (Fig. 3C; Supplemental Fig. S6A).

We observed no significant difference in the average distribution and the relative positions of tRNA between SEC-seq and Grad-seq (Unpaired t-test, P-value ≈ 0.52) (Fig. 3C). The long tail to HMW fractions that we observed in the SEC-seq data is caused by *pawZ*, a pseudogene of *argW*, which encodes tRNA-Arg^CUU^ (the fraction numbers as the relative position of *argW* and *pawZ* are 3.47 and 13.9, respectively), suggesting that *pawZ* may have molecular binding partners with which it forms HMW complexes (Supplemental Fig. S6B).

In Grad-seq, coding sequences (CDSs) are enriched in the pellet fraction, although the CDSs encoding small proteins, such as MgtS (a.k.a YneM; a regulator of the Mg^2+^ importer MgtA), co-migrated with 30S ribosomal subunits around fraction 11 (Supplemental Fig. S6C) (Hör et al. 2020a). In SEC-seq profiles, CDSs show a heterogeneous distribution with two slight peaks in fractions 11 and 19 (Fig. 3C). To investigate which factors might cause this distribution, we first performed k-means clustering of all detected CDSs according to their elution profile. The profiles were decomposed into 3 clusters based on the elbow plot (see METHOD). CDSs in cluster 3 were enriched in ribosome fractions, and the median length of the CDSs was significantly shorter than the other two clusters (one-way ANOVA with Tukey’s multiple comparison test; cluster1 = 348.8 aa, cluster 2 = 377.1 aa, and cluster 3 = 213.5 aa) (Supplemental Fig. S6E-F). This is consistent with the Grad-seq data showing co-migration of small CDS with 30S ribosomal subunits. We also performed gene ontology enrichment analysis of each cluster. Interestingly, our analysis revealed that CDSs encoding membrane proteins were enriched in cluster 1. Cluster 1 CDSs were less abundant in ribosome fractions, but showed a peak in fraction 11 (Supplemental Fig. S6E, G). Recent studies reported the translation-independent targeting of *E. coli* mRNAs encoding inner-membrane proteins to the membrane (Nevo-Dinur et al. 2011; Moffitt et al. 2016; Kannaiah et al. 2019). Our data suggest that cellular complexes might be involved in this translation-independent RNA localization. Although the SEC-seq data cannot be used to assess the subcellular localization of mRNAs, clustering analysis of highly resolved SEC-seq profiles can provide additional information about the intrinsic features of mRNAs.

Grad-seq showed enrichment of sRNAs in the pellet fraction, although higher than average abundance was also observed around fractions 3–11 (Fig. 3C). Nevertheless, this sedimentation profile mainly represents sRNAs engaged with RBPs and ribosomes to modulate translation efficiency. In contrast, sRNAs were more broadly distributed in fractions 6–20 in SEC-seq. This broader distribution likely represents sRNA-RBP complexes with different sizes and shapes, highlighting an advantage of SEC-seq in discriminating sRNA-RBP complexes as further discussed below.

### SEC-seq is able to resolve CsrA-CsrB and CsrA-CsrC complexes

A similarly broad distribution as seen for sRNAs was also observed for the RBP CsrA (Fig 4A). CsrA associates with two sRNAs antagonists, CsrB and CsrC. Since these two sRNAs have different lengths (CsrB: 369 nt vs. CsrC: 242 nt) and different numbers of CsrA-binding GGA motifs (CsrB: 18 vs. CsrC: 9) (Liu et al. 1997; Weilbacher et al. 2003), we hypothesized that the complexes they form with CsrA differ in size and shape. Although this difference is too small to detect in Grad-seq, SEC-seq is able to resolve these complexes, evidenced by the distinct elution profiles for CsrB and CsrC (Fig. 4A). To confirm that the broad distribution of CrsA in the SEC-seq profile was indeed due to the interaction with CsrB and CsrC, we constructed a C-terminally FLAG-tagged version of *csrA* in native, Δ*csrB,* Δ*csrC*, and Δ*csrBC* deletion strains and performed SEC gel-filtration followed by western blot analysis. We observed that CsrA in both native and Δ*csrC* strains peaked in fractions 15–17, while the peak shifted to fractions 13–15 in the Δ*csrB* mutant. In the Δ*csrBC* strain, CsrA was no longer detectable in HMW fractions (Fig. 4B). These results show that the elution profile of CsrA depends on its interaction with either CsrB or CsrC. In the absence of CsrB, CrsA forms a complex with CsrC, whereas in native cells the CrsA-CrsB complex dominates under our experimental conditions (Fig. 4B).

**Figure 4.**
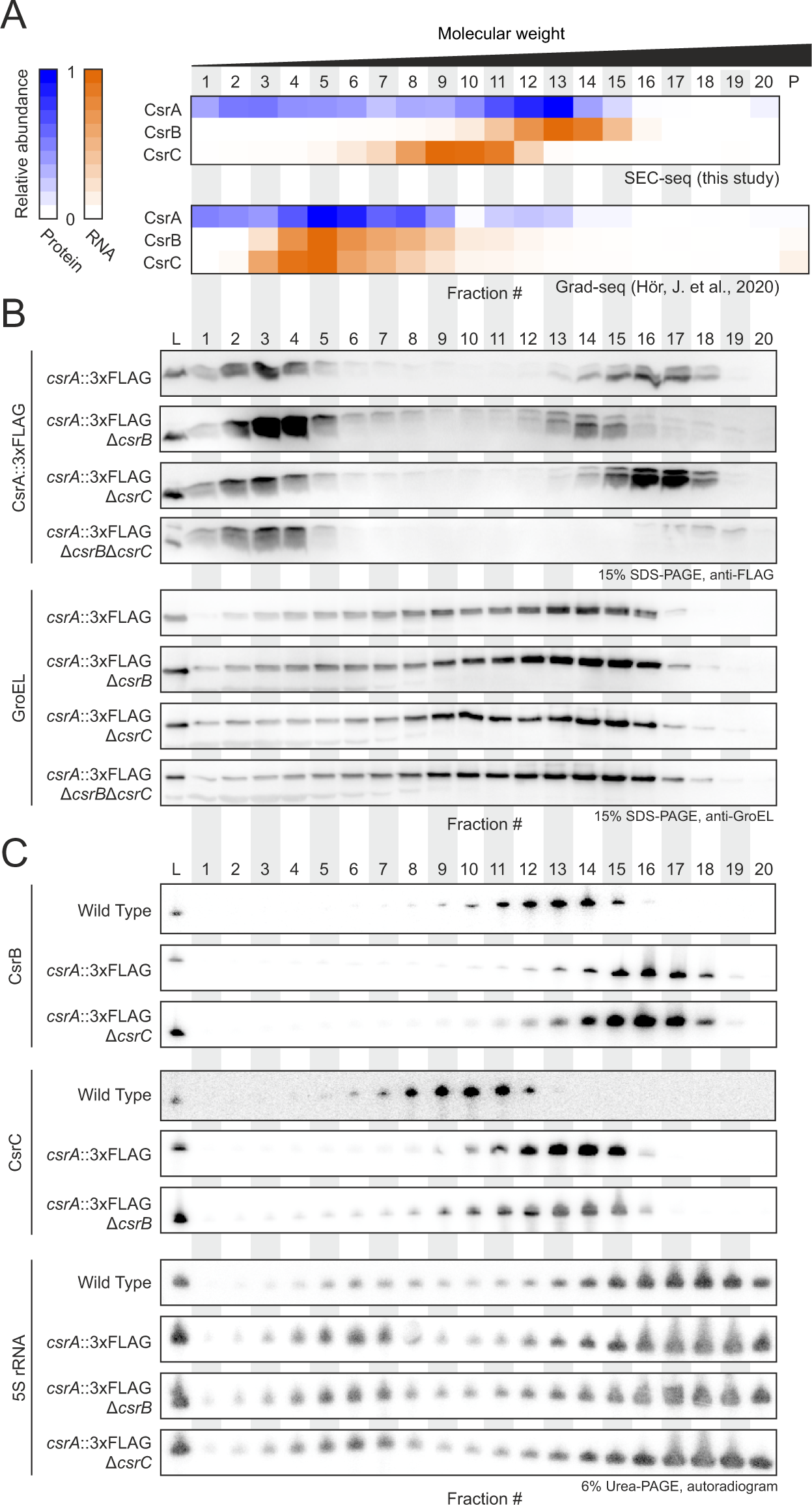
Different gel-filtration of CsrA-CsrB and CsrA-CsrC complexes in SEC. (A) Heat map of digital distribution of CsrA, CsrB and CsrC in SEC-seq and Grad-seq. The data is scaled to the maximum value. P, pellet. (B) The SEC analysis of CsrA::3xFLAG in the following mutant background strains (native, Δ*csrB,* Δ*csrC*, and Δ*csrBC*). The CsrA::3xFLAG reallocation in Δ*csrB*, Δ*csrC*, and Δ*csrBC* strains is demonstrated by western blot analysis. GroEL was used as a control. (C) The SEC analysis of CsrB/C between wild type and the *csrA::3xFLAG* strain. The CsrB/C reallocation in the *csrA::3×FLAG* strain was demonstrated by northern blot analysis. 5S rRNA is used as a control. L, lysate.

Unexpectedly, the CsrA peak in the *csrA::3×FLAG* strain was observed in fractions 15 to 17, whereas the MS data of the SEC-seq analysis of the wild type strain showed a peak in fraction 13. We speculated that the FLAG-tag itself, which is ∼2.7 kDa in size, affects the distribution of tagged CsrA. If this were the case, the positions of CsrB and CsrC would be expected to change as well. Indeed, while CsrB and CsrC peaked in fractions 13 and 10 in the wild type strain, in the *csrA::3×FLAG* strain both peaks shifted to fractions 16 and 13, respectively (Fig. 4C). In conclusion, these results demonstrate the superior ability of SEC-seq to resolve complexes within the LMW range up to 500 kDa.

### SEC-seq discriminates Hfq- and ProQ-RNA complexes

Based on prior studies, the sedimentation coefficients of Hfq- and ProQ-sRNA complexes are 11S and 5S, respectively (Smirnov et al. 2017a; Hör et al. 2020a). According to the fitted curve of the sedimentation coefficients blotted against fractions shown in Fig. 3A, both therefore sediment in similar fractions in glycerol gradients (Hfq-sRNA complexes: fraction 5; ProQ-sRNA complexes: fraction 4). Given that the molecular weight of Hfq and ProQ is relatively small (Hfq forms a hexamer of ≈ 60 kDa, ProQ is ≈ 25 kDa), we tested if SEC-seq can separate Hfq- and ProQ-binding sRNAs. We first calculated the relative peak positions of all Hfq- and ProQ-binding sRNAs that were previously validated by CLIP-seq (Holmqvist et al. 2016; Holmqvist et al. 2018), and compared their average distribution (Fig. 5A). We found that Hfq- and ProQ-binding sRNAs show approximately 5 fractions difference with respect to each median position (Kolmogorov-Smirnov test, P-value < 0.05), demonstrating the ability of SEC-seq to discriminate Hfq- and ProQ-sRNA complexes.

**Figure 5.**
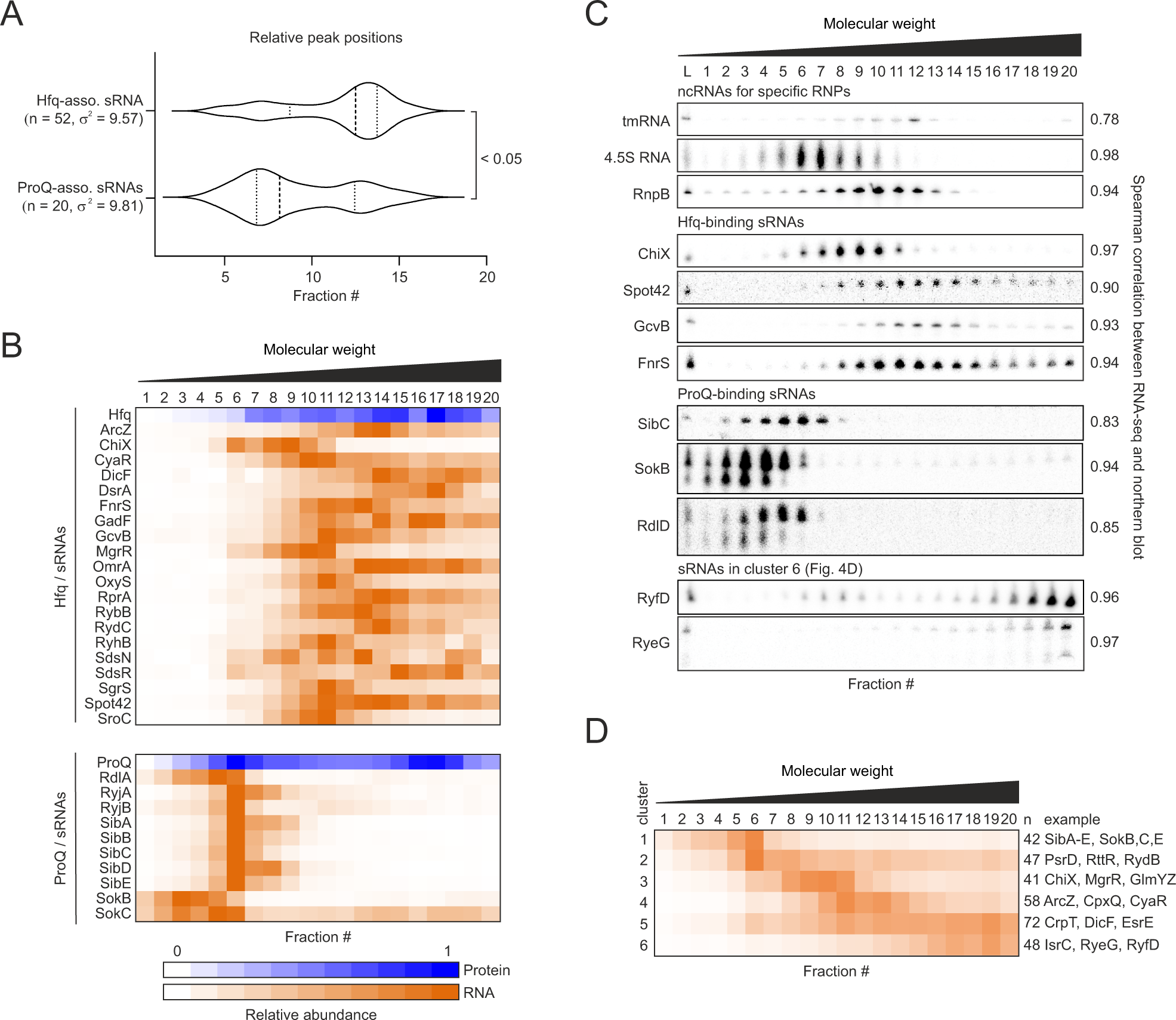
The SEC-seq profiles of Hfq- and ProQ-sRNA complexes. (A) Average of relative positions of Hfq- or ProQ-binding sRNAs in SEC-seq. All sRNAs detected by CLIP- seq with known regulatory roles are listed (Holmqvist et al. 2018; Hör et al. 2020c). Note that some sRNAs may be classified into both Hfq- and ProQ-binding sRNAs. Both Hfq- and ProQ-binding sRNAs show high variance P-value was calculated using a Kolmogorov-Smirnov test. (B) Heat map of digital distribution of representative Hfq- and ProQ- binding sRNAs. The spike-in-normalized elution profiles are normalized to the range from 0 to 1 by dividing the values of each fraction by the maximum value of the corresponding RNA and protein. (C) Northern blot analysis for the representative sRNAs distribution. Spearman’s correlation coefficient between northern blot and RNA-seq for sRNAs is shown next to each blotting result. (D) K-means clustering of all sRNAs by the elution profiles. The number of clusters were heuristically determined by elbow method. The number of sRNAs and the examples are shown next to each profile.

Next, we investigated the distribution of Hfq and ProQ and their associated sRNA in more detail (Fig. 5B) and confirmed the SEC-seq profiles of Hfq- and ProQ-binding sRNAs by northern blot analysis (Fig. 5C). Hfq was broadly distributed from fractions 7– 20 with a peak in fraction 17. This is in line with the idea that Hfq forms distinct complexes with RNAs and other proteins that differ in their size and shape (Sukhodolets and Garges 2003; Rabhi et al. 2011; Bruce et al. 2018). Many Hfq-binding sRNAs also showed a broad distribution between fractions 7 to 20, although with different peak positions. This suggests that their elution profiles depend on their association with Hfq and their various mRNA targets (Fig. 5B). To illustrate, Spot42 sRNA known to target multiple mRNAs and encode a small open reading frame (ORF) (Beisel and Storz 2011; Beisel et al. 2012; Wright et al. 2013; Aoyama et al. 2022) broadly elutes in fractions 8–20. In contrast, the sRNA ChiX elutes in fractions 6–10 (Fig. 5B). Previous reports have indicated that ChiX is destabilized through the interaction with its target mRNAs (Figueroa-Bossi et al. 2009; Overgaard et al. 2009). Thus, ChiX might not be present in HMW fractions because the Hfq-ChiX-mRNA complexes are quickly degraded. ProQ shows two distinct peaks in fractions 6 and 17 (Fig. 5B). The peaks of most ProQ-binding sRNAs correspond to fraction 6. The second ProQ peak coincides with ribosomal proteins, supporting an earlier report that ProQ associates with 30S subunits and 70S ribosomes (Sheidy and Zielke 2013).

Finally, we performed k-means clustering of all detected sRNAs according to their elution profile. The profiles fell into 6 clusters based on the elbow plot (see METHODS). The elution profiles of each cluster differed (Fig. 5D), although sRNAs in both clusters 1 and 2 peaked in fraction 6. These clusters include the known ProQ-binding sRNAs SibA- E, Sok, and PsrD. Cluster 6 was particularly interesting because some of its sRNAs exclusively localized in HMW fractions, reminiscent of the distribution of small CDSs (Fig. 5C–D). For example, cluster 6 includes the RyeG sRNA, which has previously been shown to encode a small ORF and to associate with ribosomes (Weaver et al. 2019; Hör et al. 2020a). Therefore, cluster 6 might include a group of sRNAs encoding small ORFs. Another example of a sRNA in cluster 6 is RyfD, a sRNA that lies in close proximity to *clpB,* a gene that encodes a protein disaggregase (Kawano et al. 2005). RyfD eluted in fractions 18–20, although RyfD has no obvious ORF and recent ribosome profiling approaches did not support its association with ribosomes (Meydan et al. 2019; Weaver et al. 2019). Thus, RyfD is likely to stably interact with either unknown RBPs or other RNAs. Overall, the clustering of sRNA profiles exposes similarities and differences in elution patterns and enables cross-comparison with elution profiles of potential RBPs.

### SEC analysis upon RBP deletion or RNase treatment reveals shifts in sRNA or protein distributions

One way to probe new interactions between sRNAs and potential protein partners is to examine changes in sRNA distribution after the deletion of these proteins. To apply this concept to SEC, we investigated the SEC elution profile of the known Hfq- and ProQ-binding sRNA MicA (Udekwu et al. 2005; Melamed et al. 2019) in *hfq* or *proQ* deletion strains (Fig. 6A). Northern blot analysis showed that MicA eluted in fractions 6–20 in wild type *E. coli*. In the *proQ* mutant, the peak of MicA shifted to fractions 17–20, because MicA predominantly associates with Hfq under these conditions; in the *hfq* mutant, the peak of MicA shifted to fractions 4–8 instead, indicating MicA association with ProQ. These data demonstrate the potential of SEC analysis to discover yet-unrecognized Hfq- or ProQ-binding sRNAs based on their shift in the *proQ* and *hfq* deletion strains.

**Figure 6.**
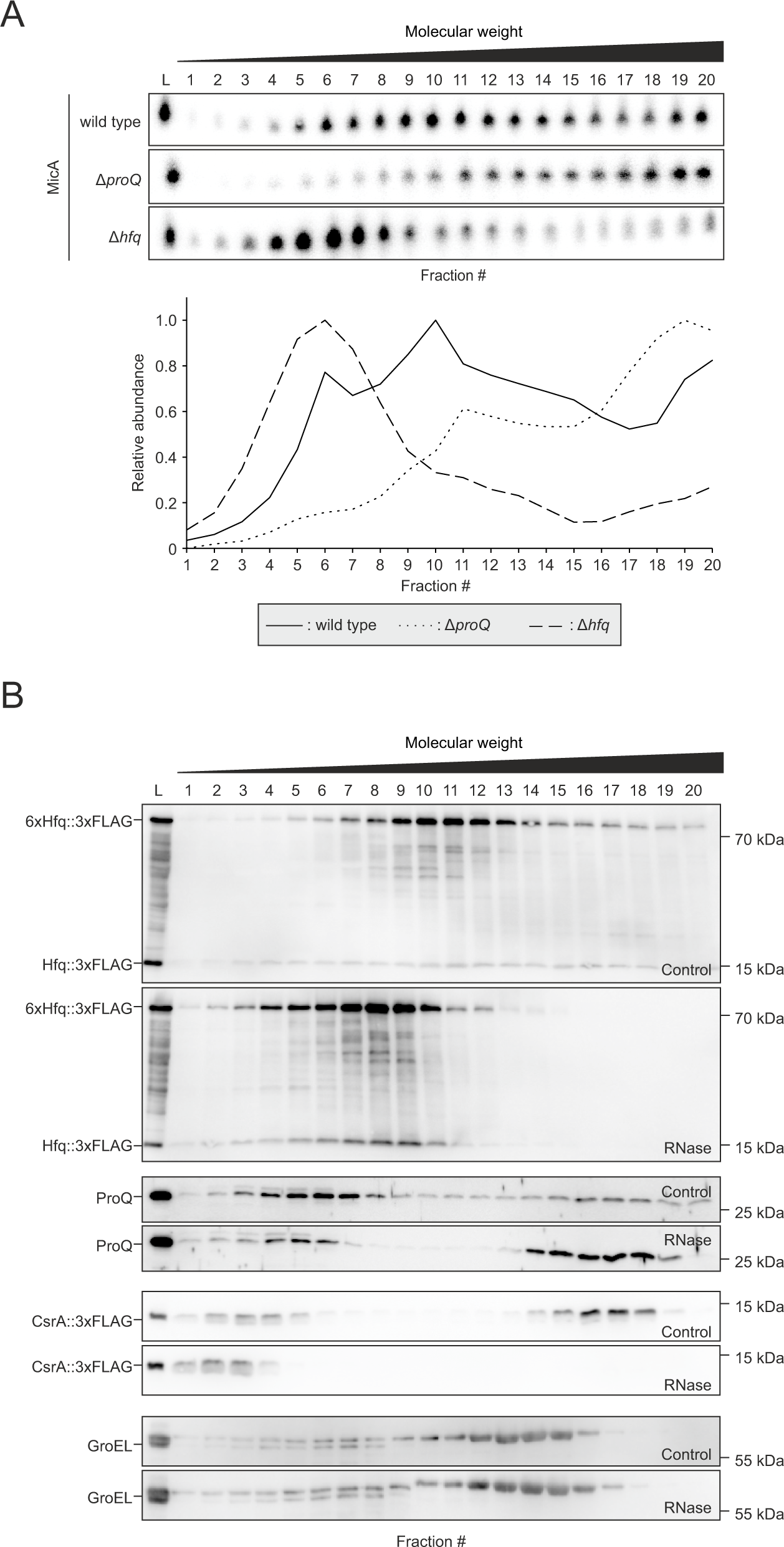
Differential SEC analysis for the interrogation of the association between RBPs and RNAs. (A) Northern blot analysis of the MicA sRNA in *E. coli* wild type, Δ*hfq*, and Δ*proQ* strains. A quantified and normalized plot is presented below the northern blot. The peak of MicA shifts to HMW and LMW fractions in the *proQ* and *hfq* deletion strains, respectively. (B) The RNase-induced shift of Hfq::3×FLAG, ProQ, CsrA::3×FLAG, and GroEL. Western blot analysis was performed for the samples with or without RNase treatment. L, lysate.

The GradR approach is based on a similar concept, as it predicts new bacterial RBPs through changes in their sedimentation profile in a glycerol gradient when associated RNAs are degraded by RNase treatment (Gerovac et al. 2020). To demonstrate that this concept can be applied to SEC, we performed SEC fractionation after RNase treatment and investigated the elution profile of the three major RBPs Hfq, ProQ, and CsrA, as well as GroEL as a negative control, by western blot analysis. After RNase treatment, the SEC chromatogram shows that peak 1, composed of aggregates, diminished, and that peak 2, which includes ribosomes, shifted to LMW fractions. This demonstrates the RNase-mediated dissociation of large aggregates and ribosomes (Supplemental Fig. S7A). Nevertheless, based on the visual inspection of SDS-PAGE gels following SEC fractionation, the distribution of the total proteome was not dramatically changed (Supplemental Fig. S7B). As expected, Hfq::3xFLAG and CsrA::3xFLAG shifted towards LMW fractions upon RNase treatment (Fig. 6B). Surprisingly, ProQ shifted toward HMW fractions upon RNase treatment. We speculate that free ProQ interacts with the Lon protease, because a previous study found that ProQ mutations that impair RNA binding stimulate its rapid turnover by Lon-mediated proteolysis (El Mouali et al. 2021). The ProQ peak in the HMW fraction indeed corresponds to the Lon peak in SEC-seq (Supplemental Table S2). Overall, our data demonstrate that SEC coupled with RNase treatment could aid the discovery of additional RBPs in bacteria.

In conclusion, SEC-seq represents a tractable and reproducible approach for the separation of bacterial RNA and protein complexes. Compared to Grad-Seq, the method shows improved resolution in the LMW range, a size range that is particularly relevant for the analysis of regulatory RNA-protein complexes in bacteria. Therefore, SEC-seq has the potential to become the gold-standard for the prediction of bacterial RNA-protein complexes.

## METHODS

### Bacteria and growth media

*E. coli* K-12 MG1655, *P. aeruginosa* PAO1 and the derivatives were streaked on Luria-Bertani (LB) agar plates and grown overnight at 37°C. Overnight cultures were prepared at 37°C in LB medium with shaking at 220 rpm. *C. difficile* was grown in Brain Heart Infusion broth under anaerobic conditions inside a Coy chamber (85% N_2_, 10% H_2_, and 5% CO_2_). Antibiotics were used as needed at concentrations listed as follows: 100 μg/ml carbenicillin and 50 μg/ml kanamycin.

### Strain construction

All strains, plasmids, and oligonucleotides are listed in Supplemental Tables S4, respectively. Deletion and 3×FLAG tagged mutants were generated as previously described (Datsenko and Wanner 2000; Uzzau et al. 2001). Briefly, the overnight culture of *E. coli* MG1655 with pKD46 was 100-fold diluted into 50 mL of fresh LB medium containing 0.2% arabinose and grown to OD_600_ = 0.5 at 28°C. Thereafter, bacteria are pelleted by centrifugation at 4°C at 4,100 ×g for 20 min, and washed with ice-cold 10% glycerol twice. Cells were concentrated at 500 μl, and 800 ng of PCR product was added to 80 μl of obtained electrocompetent cells. pKD4 and pSUB11 plasmids were used as a temperate for PCR amplification of the kanamycin gene cassette and 3×FLAG, respectively (Datsenko and Wanner 2000; Uzzau et al. 2001). The transformants were streaked on LB agar with 50 μg/ml kanamycin and incubated overnight. P1 phage lysates were prepared from obtained mutants and transduced into *E. coli* MG1655 wild type strain (Thomason et al. 2007). For sequential mutations, the mutants were cured by pCP20 carrying the FLP recombinase as previously described (Datsenko and Wanner 2000).

### Fractionation by size exclusion chromatography

Overnight cultures of *E. coli* MG1655, *Salmonella* SL1344, *P. aeruginosa* PAO1 or their derivatives were 100-fold diluted in 100 ml of fresh LB medium, grown to an OD_600_ of 2.0, cooled down in an ice-water bath for 15 min, and then harvested by centrifugation for 20 min at 4°C and 4,100 ×g. Overnight culture of *C. difficile* 630 was 100-fold diluted in 20 ml of fresh BHI medium and grown for 4 h. Thereafter, the pre-culture was again 100- fold diluted in 200 ml of fresh BHI medium, grown to an OD_600_ of 1.0, cooled down in an ice-water bath for 15 min, and then harvested as mentioned above. The cells were washed three times with ice-cold 1× TBS [20 mM Tris, 150 mM NaCl, pH 7.6], resuspended in 500 μl of ice-cold 1× lysis buffer [20 mM Tris–HCl, pH 7.5, 150 mM KCl, 1 mM MgCl_2_ (10 mM MgCl_2_ for *Salmonella*), 1 mM DTT, 1 mM PMSF, 0.2% Triton X 100, 20 U/ml DNase I (Thermo Fisher, cat#EN0521), 200 U/ml RNase inhibitor] and lysed by addition of 1× volume of 0.1 mm glass beads and 10 cycles of vortexing for 30 s followed by cooling on ice for 15 s. To remove insoluble debris and the glass beads, the lysate was cleared by centrifugation for 10 min at 4°C and 16,100 ×g. Ten microliters of cleared lysate were mixed with 1 ml TRIzol (Thermo Fisher, cat# 15596026) for the RNA input control, and 20 μl was mixed with 20 μl 5× protein loading buffer for the protein input control. Before loading onto the column, Superose 6 Increase 10/300 GL column (GE Healthcare) was equilibrated with the twice volume of 1× lysis buffer without PMSF, DNase I, and RNase Inhibitor. The cleared lysate was then injected into the equilibrated column connected to ÄKTA pure purification system (GE Healthcare). We did not observed the clogging of the column and the system. The flow rate was set to 0.25 ml/min, and the elution volume was set to 0.38 ml. Each fraction was automatically collected using fraction collector F9-R (GE Healthcare). Fractionation was operated at 4°C. For the experiments with RNase treatment, 100 μl of RNase A/T1 mix (2 μg/μl, 5 U/μl, Thermo Scientific, cat#EN0551) was added to 400 μl of the cleared lysate and incubated for 20 min at room temperature. The reaction was stopped on ice and loaded into the equilibrated column connected to ÄKTA pure purification system (GE Healthcare) as described above. For protein analysis, 90 μl of each fraction was taken and mixed with 30 μl of 5× protein loading buffer. The remaining 290 μl of each fraction was used for RNA isolation by 25 μl of 10% SDS and 1× volume of acidic phenol/chloroform/isoamylalcohol (P/C/I). The fractions were then vortexed for 30 s and let rest at room temperature for 5 min before separating the phases by centrifugation for 15 min at 4°C and 16,100 ×g. The aqueous phases were precipitated with 1 μl of GlycoBlue (Thermo Fisher, cat#AM9515) and 2× volume of ice-cold ethanol/3M sodium acetate, pH 6.5 (30:1) overnight at −20°C. The RNA was collected by centrifugation for 30 min at 4°C and 16,100 ×g and washed with 1× volume of ice-cold 70% ethanol, followed by centrifugation for 15 min at 4°C at 16,100 ×g. The lysate RNA sample stored in TRIzol was purified according to the manufacturer’s protocol, except that the precipitation was performed using the ethanol mix mentioned above. After drying of the RNA pellet, it was dissolved in 40 μl DEPC-treated H_2_O and DNase-digested by the addition of 5 μl DNase I buffer with MgCl_2_, 0.5 μl RNase inhibitor, 4 μl DNase I, and 0.5 μl DEPC-treated H_2_O, followed by incubation for 45 min at 37°C. The DNase-treated RNA was purified by adding 100 μl DEPC-treated H_2_O and 1× volume of P/C/I as described above. The purified, DNase-treated RNA was dissolved in 35 μl DEPC-treated H_2_O.

### RNA gel electrophoresis and northern blotting

Equal volumes of the extracted RNAs from each 20 fraction were separated by 6% denaturing PAGE in 1× TBE and 7 M urea and stained with ethidium bromide. For northern blotting, unstained gels were transferred to Hybond+ membranes (GE Healthcare) followed by UV crosslink with 120 mJ of UV light at 254 nm. The UV crosslinked membrane was probed with RNA-specific radioactively labeled DNA oligonucleotides in ROTI^®^Hybri-Quick (CARL Roth, cat#A981.2) at 42°C overnight. Probed membranes were washed every 15 min in 5× Saline Sodium Citrate (SSC)/0.1% SDS, 1× SSC/0.1% SDS and 0.5× SSC/0.1% SDS buffers at 42°C. Autoradiograms were visualized with Typhoon FLA 7000 (GE Healthcare) and quantified using ImageJ.

### Protein gel electrophoresis and western blotting

Equal volumes of the protein samples from each 20 fraction were separated by 12% or 15% SDS-PAGE and stained with ROTI^®^-Blue (CARL Roth, cat#A152.1) according to the manufacturer’s protocol. For western blotting, unstained gels were transferred to PVDF membranes (GE Healthcare) and probed with protein-specific antisera against RpoB (1:10,000 dilution, BioLegend, cat# 663905), 3xFLAG (1:1,000 dilution, Sigma-Aldrich, cat#F1804), ProQ (1:10,000 dilution, kind gift of Daniel Sheidy), or GroEL (1:1,000 dilution, Sigma-Aldrich, cat#G6532) diluted in 1× TBS-T buffer [20 mM Tris, 150 mM NaCl, 0.1% Tween20, pH 7.6] containing 3% bovine serum albumin. After washing with 1× TBS-T buffer three times, membranes were probed with anti-mouse-HRP-antibody (1:10,000 dilution, Thermo Fisher, cat# 31430) or anti-rabbit-HRP-antibody (1:10,000 dilution, Thermo Fisher, cat# 31460). Chemiluminescent signals were visualized with ECL™ Select western blotting detection reagent (Cytiva, cat#RPN2235) and measured with Image Quant LAS 4000 (GE Healthcare).

### RNA-seq

RNA-seq was performed as described before (Hör et al. 2020a) at Vertis Biotechnologie AG. Briefly, 5 μl of the purified RNAs were 10-fold diluted in DEPC-treated H_2_O. Ten- micro-liter of the aliquot was mixed with 10 μl of a 1:100 dilution of the ERCC spike-in mix 2 (Thermo Fisher). The resulting RNA samples were fragmented, ligated with 3’ adapter, and reverse-transcribed with MMLV reverse transcriptase. Purified first-strand cDNA was then ligated with the 5’ Illumina TruSeq sequencing adapter, followed by PCR amplification to about 10-20 ng/μl using a high-fidelity DNA polymerase. The cDNA samples were purified with the Agencourt AMPure XP kit (Beckman Coulter) and pooled with ratios according to the input samples’ RNA concentrations. Finally, cDNA with a size range of 200-550 bp was gel-eluted using a preparative agarose gel. The pooled libraries were sequenced on an Illumina NextSeq 500 system, and 75 nt single-end reads were generated.

### RNA-seq data analysis

Read trimming and clipping were done with cutadapt (Martin 2011). Read filtering, read mapping, nucleotide-wise coverage calculation, and genome feature-wise read quantification were done using READemption (Forstner et al. 2014) (v0.4.3) and the short-read mapper segemehl v0.2.0 (Hoffmann et al. 2014) using the *Escherichia coli* MG1655 genome (accession number NC_000913.3) as reference. The annotation provided was extended by ncRNAs predicted by ANNOgesic (Yu et al. 2018). The analysis was performed with the tool GRADitude (Di Giorgio, S., Hör, J., Vogel, J., Förstner, K.U., unpublished; v0.1.0; https://foerstner-lab.github.io/GRADitude/). Only transcripts with a sum of ≥ 100 reads in all fractions were considered for the downstream analysis. Read counts for each fraction were normalized by calculating size factors following the DESeq2 approach (Love et al. 2014) generated from the ERCC spike-in read counts added to each sample (see above). To make all the transcript counts comparable, they were scaled to the maximum value. After normalization, k-means clustering (Lloyd 1982) and t-SNE (T-distributed stochastic neighbor embedding) dimension reduction (van der Maaten and Hinton 2008) was performed using the Python package scikit-learn. All default parameters provided by the sklearn.manifold.TSNE class were used.

### Sample preparation for mass spectrometry

Sample preparation for mass spectrometry was performed as described before (Hör et al. 2020a). Briefly, the protein samples (diluted in 1.25× protein loading buffer) were sonicated with 5 cycles of 30 s on followed by 30 s off at 4°C (Bioruptor Plus, Diagenode). After the centrifugation for 15 min at 4°C and 16,100 ×g, 20 μl of the soluble protein sample were mixed with 10 μl of UPS2 spike-in (Sigma-Aldrich, cat# UPS2) diluted in 250 μl 1.25× protein loading buffer. Reductive alkylation was performed by incubation with 50 mM DTT for 10 min at 70°C followed by incubation with 120 mM iodoacetamide for 20 min at room temperature in the dark. After the precipitation with four volumes of acetone overnight at -20°C, pellets were dissolved in 50 μl of 8 M urea with 100 mM ammonium bicarbonate. The precipitated proteins were then digested with 0.25 μg Lys- C (Wako) for 2 h at 30°C, followed by digestion with 0.25 μg trypsin overnight at 37°C. The digested peptides were desalted with 60% acetonitrile/0.3% formic acid through the three disks of C-18 Empore SPE disks (3M) in a 200 μl pipet tip. After drying in a laboratory freeze-dryer (Christ), the peptides were dissolved in 2% acetonitrile/0.1% formic acid.

### Nano LC-MS/MS analysis

NanoLC-MS/MS analysis was performed as described before (Hör et al. 2020a) at Rudolf-Virchow-Center Würzburg for Integrative and Translational Bioimaging. The peptides were loaded on capillary columns (PicoFrit, 30 cm × 150 μm ID, New Objective) filled with ReproSil-Pur 120 C18-AQ, 1.9 μm (Dr. Maisch), which is connected with an Orbitrap Fusion (Thermo Scientific) equipped with a PicoView Ion Source (New Objective) and an EASY-nLC 1000 liquid chromatography system(Thermo Scientific). Peptides were then separated with a 140 min linear gradient from 3% to 40% acetonitrile and 0.1% formic acid at a flow rate of 500 nl/min. Both MS and MS/MS scans were acquired in the Orbitrap analyzer with a resolution of 60,000 for MS scans and 15,000 for MS/MS scans. HCD fragmentation with 35% normalized collision energy was applied. A Top Speed data-dependent MS/MS method with a fixed cycle time of 3 s was used. Dynamic exclusion was applied with a repeat count of 1 and an exclusion duration of 60 s; singly charged precursors were excluded from selection. The minimum signal threshold for precursor selection was set to 50,000. Predictive AGC was used with a target value of 2×10^5^ for MS scans and 5×10^4^ for MS/MS scans. EASY-IC was used for internal calibration.

### MS data analysis

Raw MS data files were analyzed with MaxQuant version 1.5.7.4 (Tyanova et al. 2016). The search was performed against the UniProt database for *E. coli* MG1655 (organism identifier: ECOLI), UPS2 spike-in, and common contaminants. The search was conducted with tryptic cleavage specificity with 3 allowed miscleavages. Protein identification was under the control of a false-discovery rate of 1% on both protein and peptide levels. In addition to the MaxQuant default settings, the search was performed against the following variable modifications: Protein N-terminal acetylation, Gln to pyro-Glu formation (N-terminal Q), and oxidation on Met. For protein quantitation, the LFQ intensities were used (Tyanova et al. 2016). Proteins with less than 2 identified razor/unique peptides were dismissed. Normalization of the proteins across the fractions was performed using the UPS2 spike-in. For this, only spike-in proteins with detectable intensities in all fractions were used. The spike-in proteins showing the highest variance (median average deviation of log_10_ intensities >1.5x lQR) were eliminated. Following this, for each spike-in protein, the median log_10_ intensity was subtracted from each fraction’s log_10_ intensities. The fraction-wise median of the resulting values was then removed from the log_10_ intensities for each bacterial protein in the corresponding fractions. Finally, all log_10_ intensities smaller than the 5% quantile of all intensities in the data set were replaced by the 5% quantile value of all intensities in the data set.

### Statistical and other analyses

The localization of SEC-detected proteins was searched in EcoCyc (v. 22.0) (Keseler et al. 2017). A Smarttables was created based on the type “complexes, proteins.” Redundant proteins across the different complexes were removed.

Gene ontology analysis was performed using DAVID 6.8 (Huang et al. 2009). Enrichment was calculated based on the identifier “UNIPROT_ACCESSION.” EASE value was set to 0.01. Among the annotation summary results, Gene Ontology term “cellular compartment” was used throughout the manuscript. All results including GO terms “biological process” and “molecular function” are shown in Supplemental Table S3.

The mean position of RNA in fractions was calculated as follows. First, the relative abundance of RNA normalized to the range from 0 to 1 was multiplied by the number of the corresponding fraction. Second, the values from all fractions (fractions 1–20) were summed up and divided by 20 which is the total fraction number. This value is regarded as the mean position of RNA in fractions.

Microsoft Excel was used to produce a heat map of the average RNA and protein distributions in SEC (e.g., Fig. 1E) and the average and the standard deviation of relative abundance of RNA (e.g., Fig. 2C). GraphPad Prism 9.1.0 was used to calculate fraction-wise and gene-wise Pearson correlation of RNA and protein distributions (Supplemental Fig. S5), to perform an unpaired *t*-test against the mean position of each RNA class and visualize them as violin plot (e.g., Fig. 5A), and to analyze spearman correlation of the RNA distribution between northern blot and RNA-seq (Fig. 5C).

## DATA DEPOSITION

The Sequencing data have been deposited in NCBI’s GENE Expression Omnibus (Edgar et al. 2002), and are accessible through GEO Series accession number GSE212408 (https://www.ncbi.nlm.nih.gov/geo/query/acc.cgi?acc=GSE212408). The mass spectrometry proteomics data have been deposited to the ProteomeXchange Consortium (Deutsch et al. 2020) via the PRIDE partner repository (Perez-Riverol et al. 2019) with the dataset identified PXD036475 (https://www.ebi.ac.uk/pride/archive/projects/PXD036475). The RNA and protein elution profiles can be viewed through an open access online browser under https://helmholtz-hiri.de/en/datasets/gradseqec/. READemption 0.4.3 is deposited at Zenodo 250598 (https://zenodo.org/record/250598).

## ACKNOWLEDGEMENTS

We thank Anke Sparmann for editing the manuscript and Svetlana Durica-Mitic, Falk Ponath, and other members of the Vogel lab for comments and fruitful discussions. We thank Andreas Schlosser and Stephanie Lamer from the Rudolf-Virchow-Center, Würzburg, for mass spectrometry analysis; Silvia Di Giorgio (supervised by Konrad Förstner) for executing the READemption and GRADitude scripts on one of the replicates of RNA-seq data and for normalizing one replicate of the MS data; Tina Lence for technical help with preparing the *C. difficil*e cultures. This work was funded by a Gottfried Wilhelm Leibniz award to J.V. (DFG grant 875-18).

## Authors’ contributions

K.C., J.H., and J.V. designed the experiments. K.C. conducted the experiments, performed computational analyses, and generated all figures. M.G. programmed the SEC-seq browser. K.C. and J.V. wrote the manuscript with input from all authors.

## FIGURE LEGENDS

**Supplemental Figure S1.**
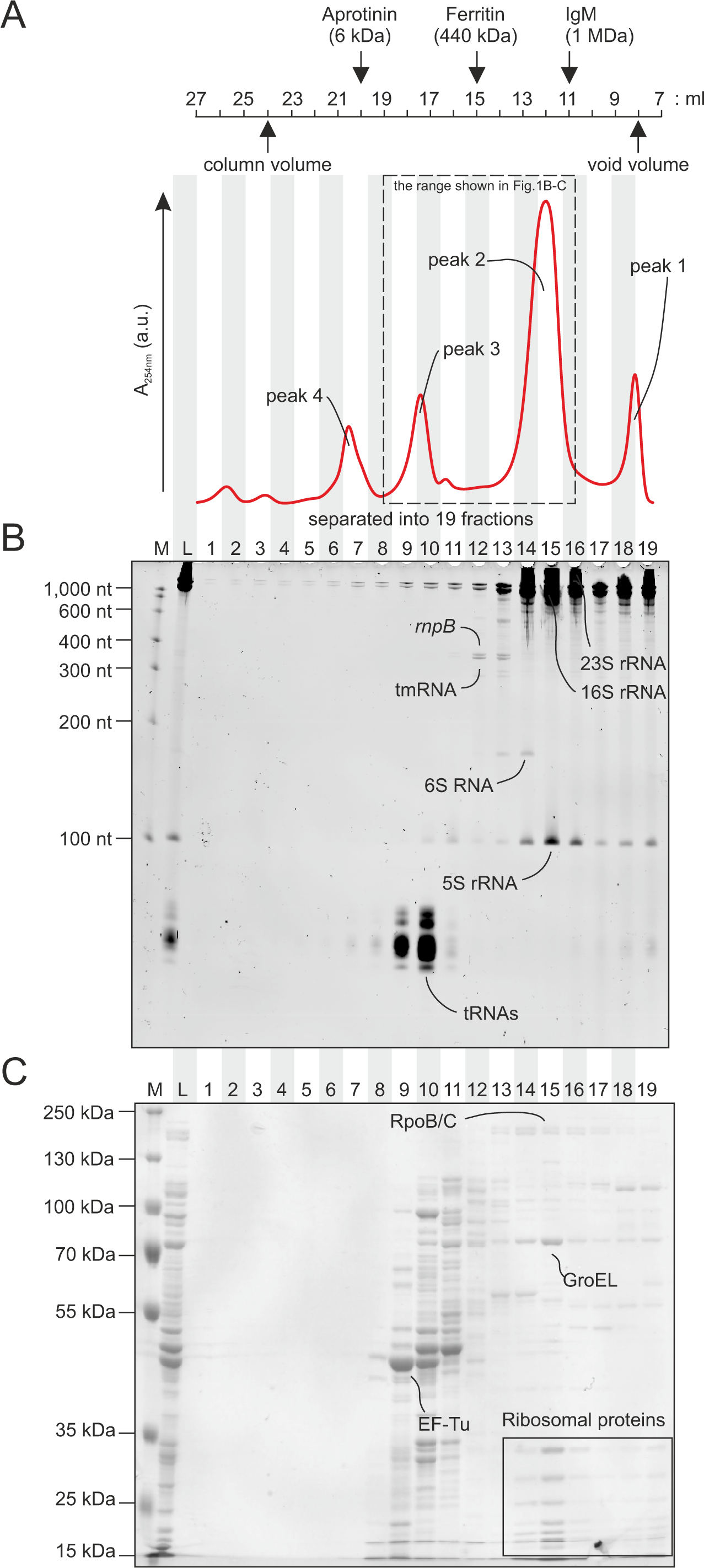
Preliminary investigation of the elution profile of RNAs and proteins. (A) The SEC chromatogram at 280 nm. The peaks of aprotinin, ferritin, and IgM are shown as standard proteins from the manual of Superose 6 Increase 10/300 GL (Cytiva 2020). (B and C) Visualization of RNAs (B) and proteins (C) elution profiles by Urea-PAGE and SDS-PAGE. High abundant house-keeping RNAs and proteins are indicated. M, marker. L, lysate.

**Supplemental Figure S2.**
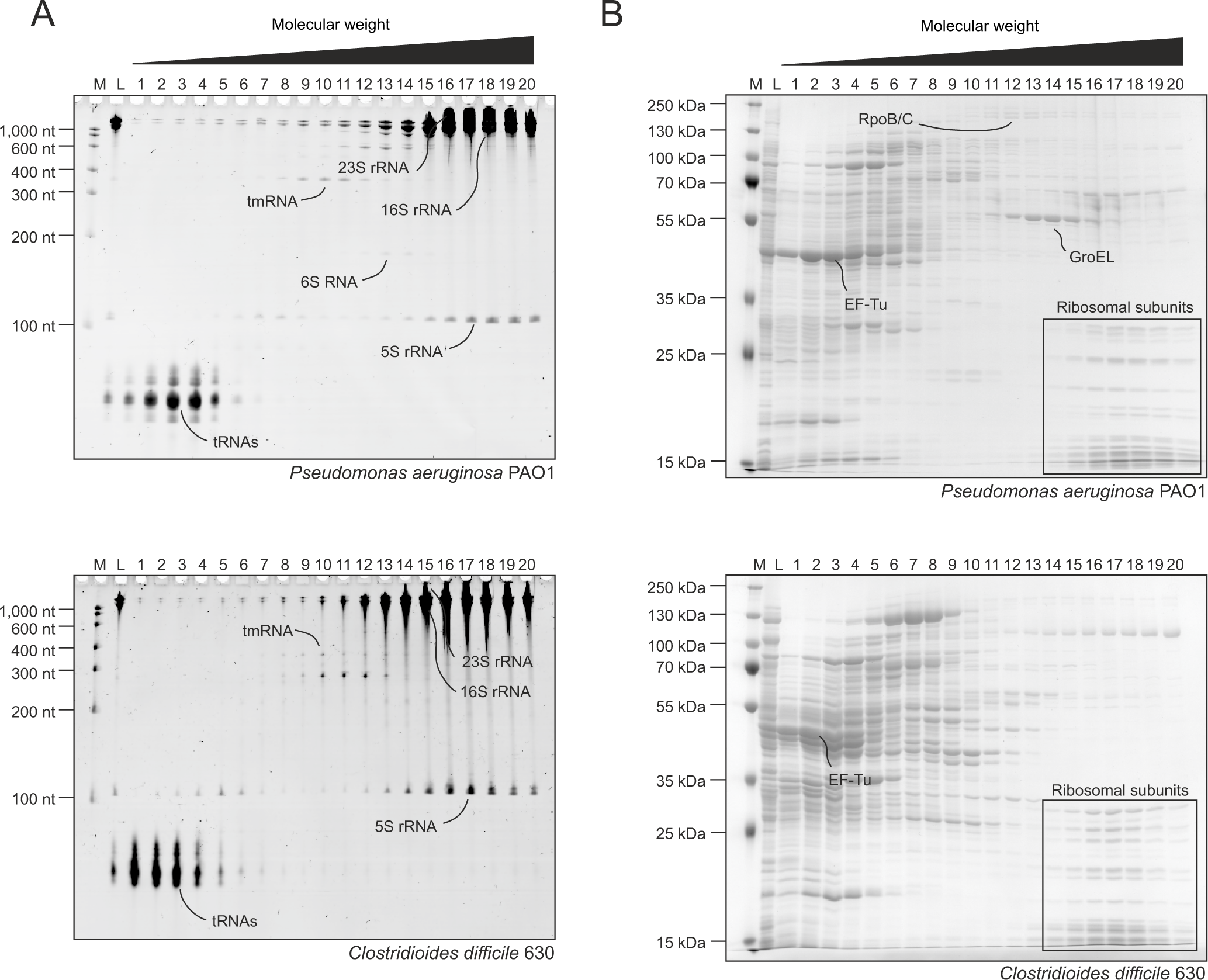
The SEC profile of RNAs and proteins in *P. aeruginosa* and *C. difficile*. (A and B) The SEC elution profiles of RNAs (A) and proteins (B) visualized by Urea-PAGE and SDS-PAGE. The overall profiles of representative RNAs and proteins are similar with *E. coli*.

**Supplemental Figure S3.**
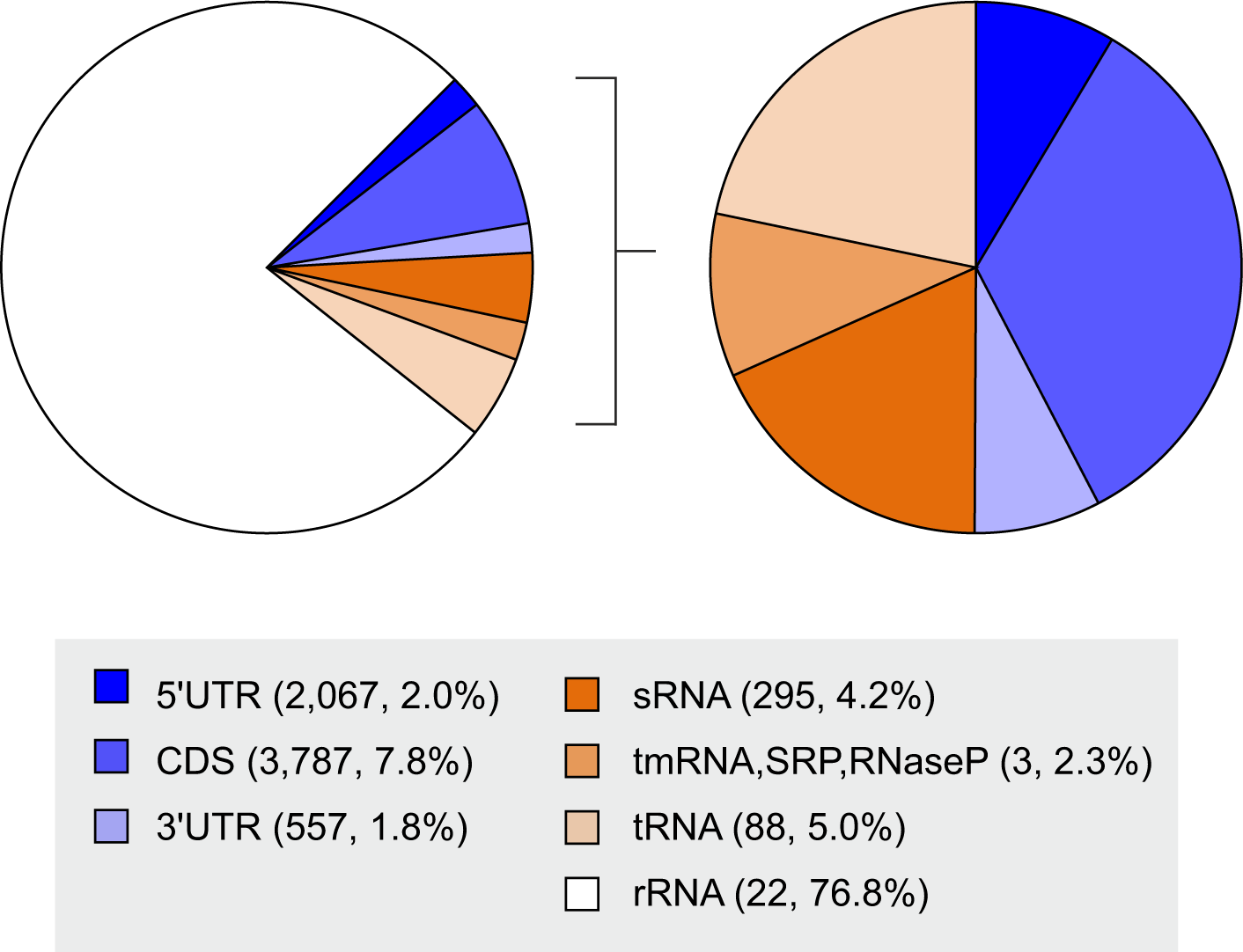
Abundance of RNA detected by SEC-seq. Ratio of reads of each RNA class detected by SEC-seq. Approximately three-quarters of reads come from rRNA. sRNAs including tmRNA, SRP, and RNase P contributes 6.5% of all reads.

**Supplemental Figure S4.**
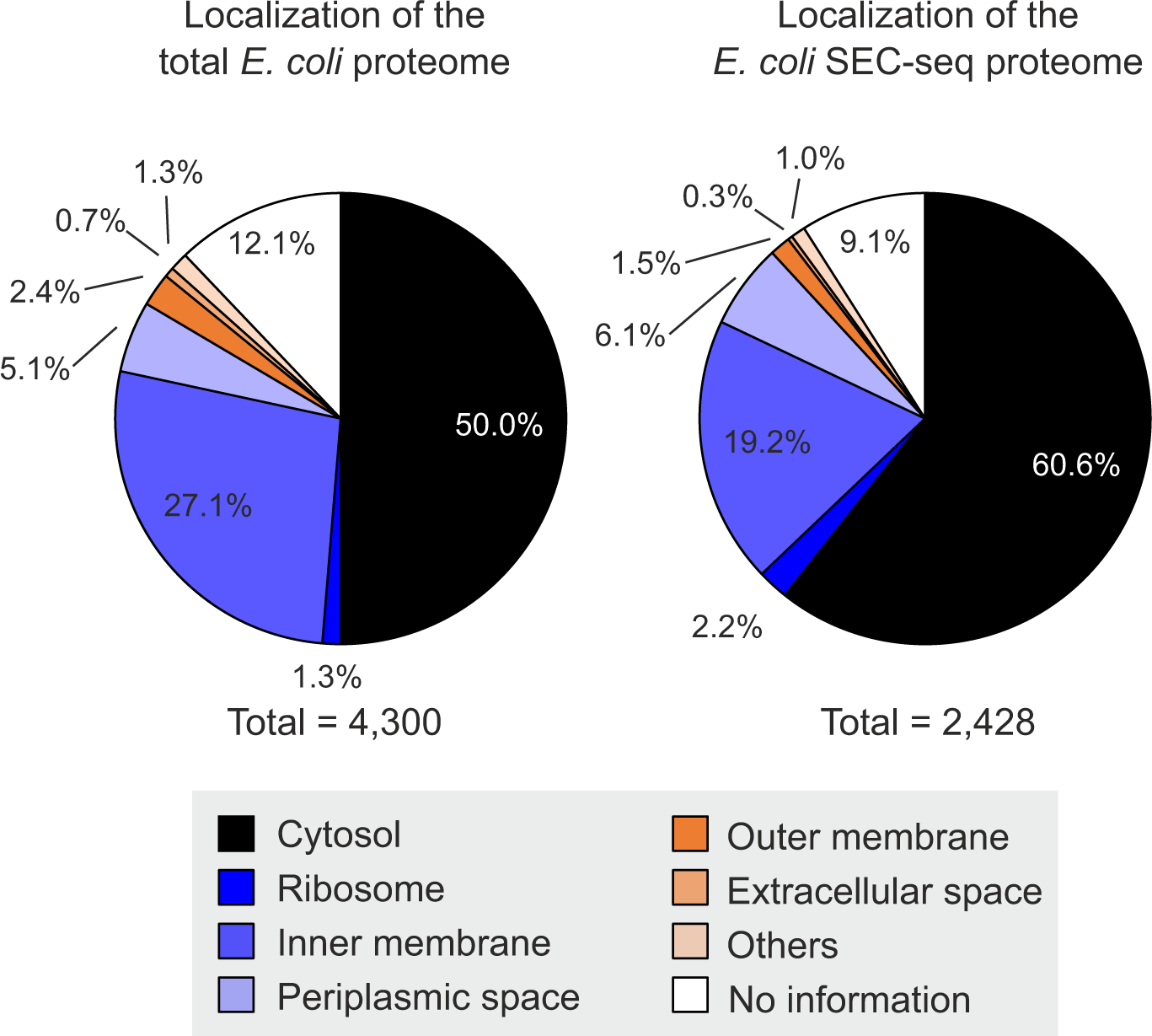
Localization of proteins detected by SEC-seq. Localizations of all *E. coli* proteins reported by EcoCyc and the proteins detected by SEC-seq are shown. Note that some proteins have more than one localization assigned. The *E. coli* SEC-seq primarily recovered cytosolic proteins constituting 60% of the ∼2,400 detected proteins.

**Supplemental Figure S5.**
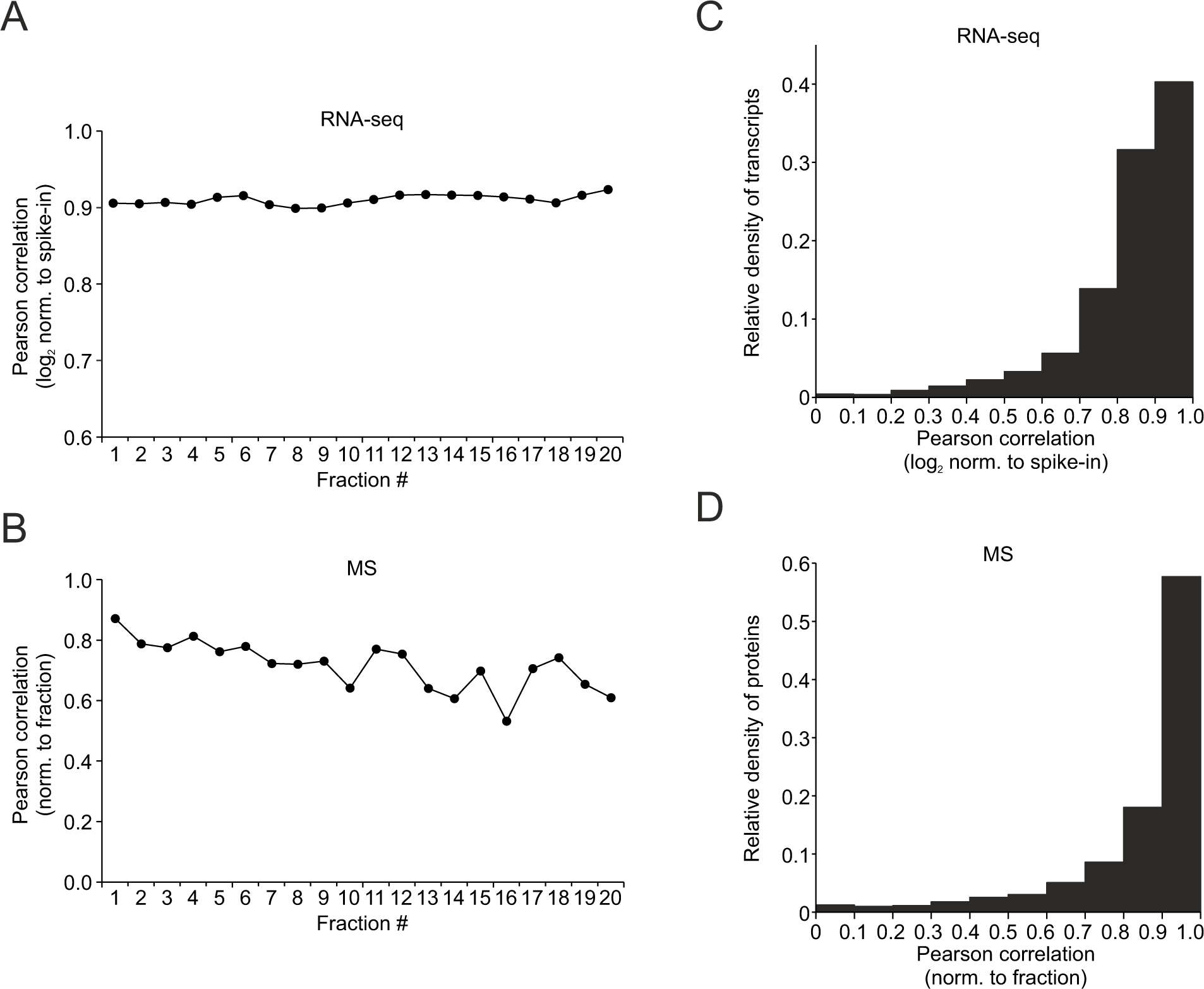
The reproducibility of SEC-seq. (A and B) Fraction-wise Pearson correlation of RNAs (A) and proteins (B) between two independent biological replicates. Content and abundance of all detected RNAs and proteins in each fraction were compared between replicates. (C and D) Gene-wise Pearson correlation of RNAs (C) and proteins (D) between two independent biological replicates. The normalized distributions in each RNA and protein were compared and the calculated Pearson correlation coefficient is shown as cumulative plots.

**Supplemental Figure S6.**
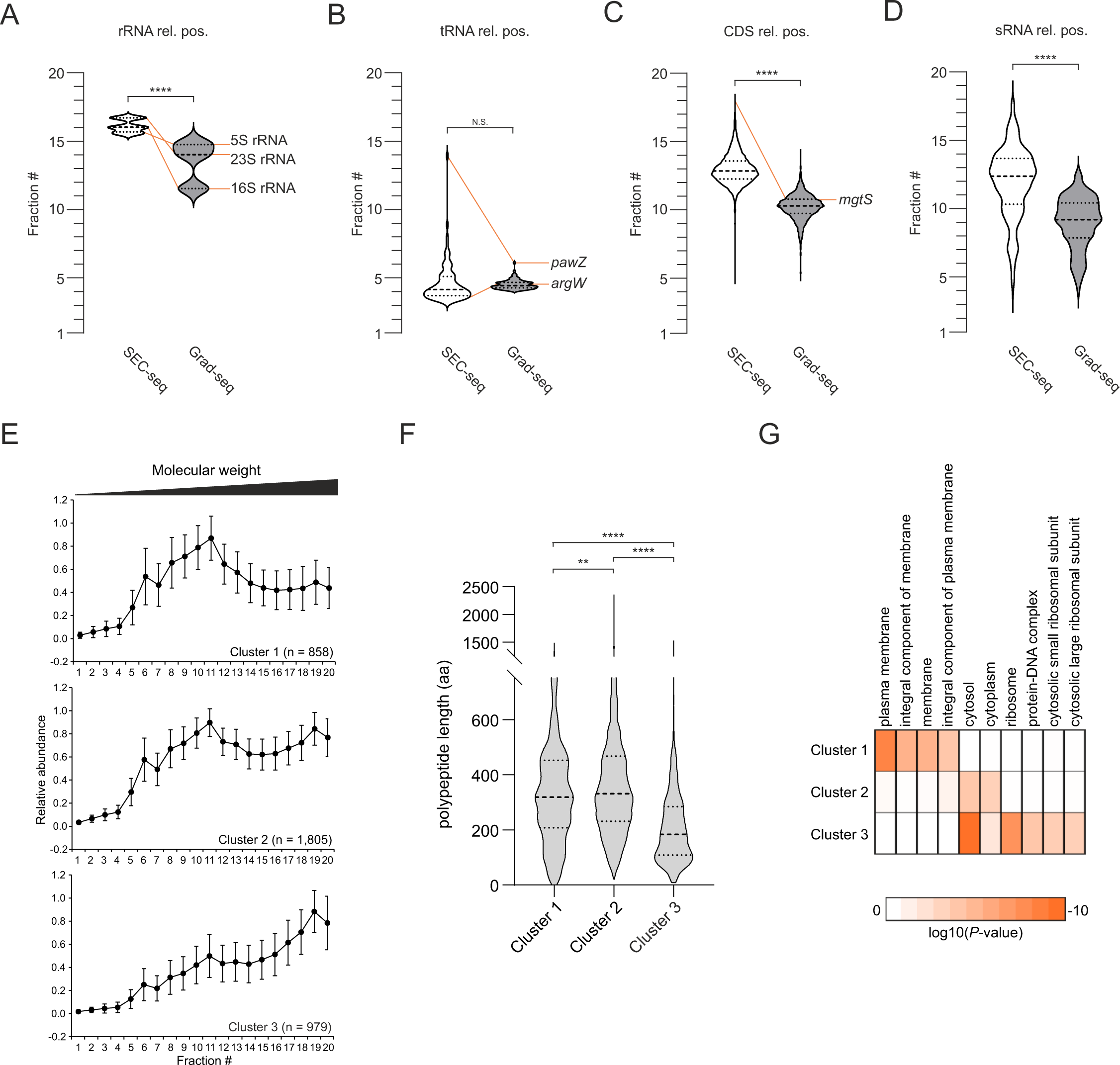
Average of relative positions of RNAs. (A–D) Average of relative positions is shown for rRNA (A), tRNA (B), CDS (C), and sRNA (D). Unpaired t-test is performed against the relative position in each RNA class (rRNA, tRNA, CDS, and sRNA): ****, p-value < 0.0001; N.S., not significant. Note that the pellet fraction is removed in Grad-seq. (E) Decomposition of the SEC-seq profile for all detectable mRNAs into three clusters. The average and the standard deviation of relative abundance scaled to the maximum values are calculated. (F) Violin plot of the distribution of mRNA length in each cluster. The bold dashed line indicates the median, whereas the dotted lines indicate the upper and lower quartiles. Statistical analyses are performed using one-way ANOVA with Tukey’s multiple comparison test): ****, p-value < 0.0001; **, p-value < 0.005. (G) DAVID enrichment analysis of mRNAs in each cluster (Huang et al. 2007). The terms with Fisher’s exact *P*-value lower than 0.01 in GO term “cellular component” are presented. All results including GO terms “biological process” and “molecular function” are shown in Supplemental Table S3.

**Supplemental Figure S7.**
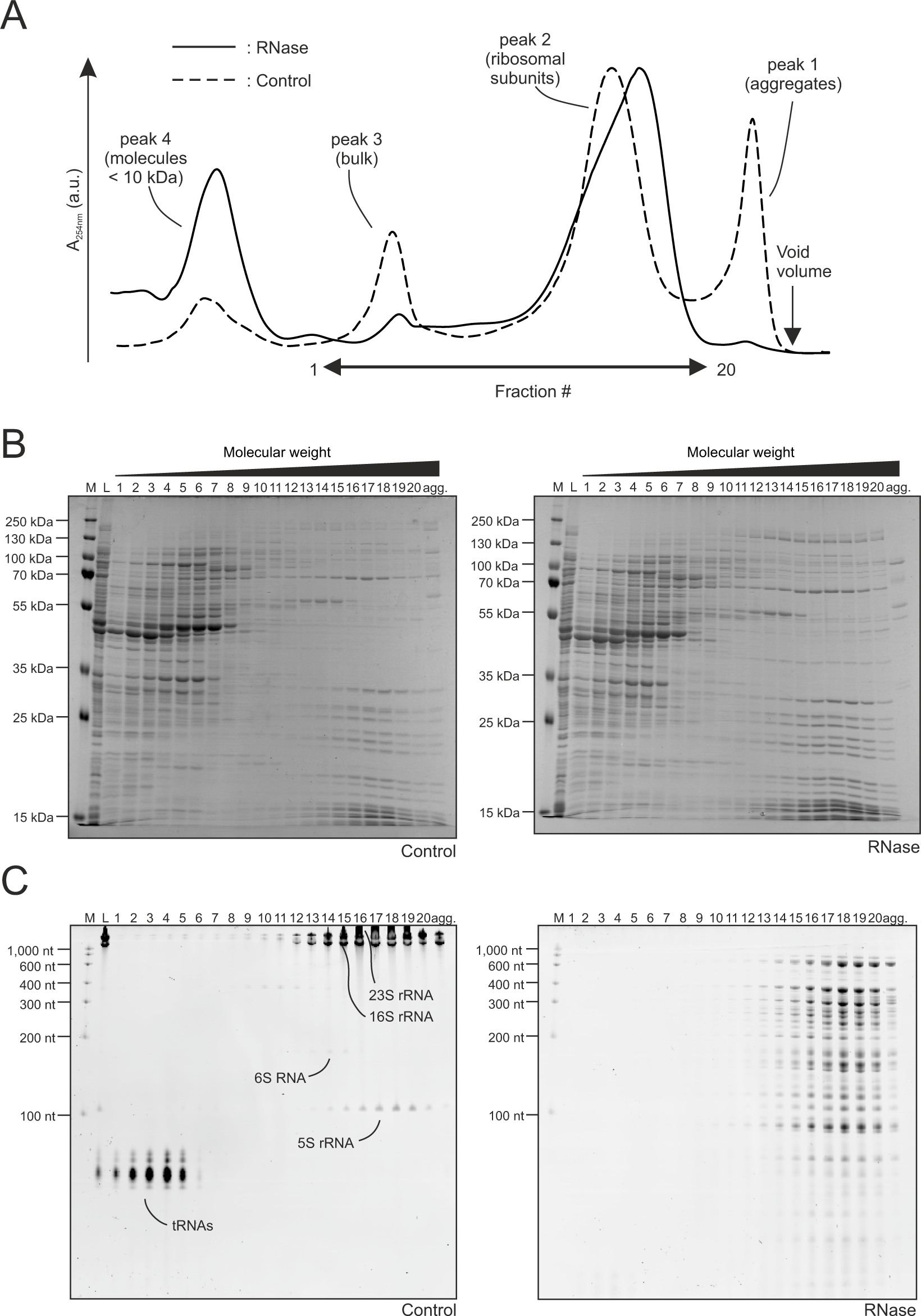
The SEC-seq profiles of RNAs and proteins with RNase treatment. (A) Comparison of chromatograms with or without RNase treatment. The peaks from polysomes and bulk decreased in the RNase treatment. In contrast, the peak probably derived from degradation products increased in the RNase treatment. (B and C) Visualization of different proteins (B) and RNAs (C) migration with or without RNase treatment. The allocation of the total proteome is not changed: agg., pool of fractions from aggregates. M, marker. L, lysate.

**Supplemental Table S1 SEC-seq dataset**

**Supplemental Table S2 SEC-MS dataset**

**Supplemental Table S3 DAVID enrichment analysis result**

**Supplemental Table S4 Strains, plasmids, and oligonucleotides used in this study**

